# Rapid and efficient generation of antigen-specific isogenic T cells from cryopreserved blood samples

**DOI:** 10.1101/2021.11.12.468355

**Authors:** AL Eerkens, A Vledder, N van Rooij, F Foijer, HW Nijman, M de Bruyn

**Affiliations:** University of Groningen, University Medical Center Groningen, Department of Obstetrics and Gynecology, The Netherlands; University of Groningen, University Medical Center Groningen, European Research Institute for the Biology of Ageing, The Netherlands

## Abstract

**Objectives:** CRISPR/Cas9-mediated gene editing has been leveraged for the modification of human and mouse T cells. However, limited experience is available on the application of CRISPR/Cas9 electroporation in cryopreserved T cells collected during e.g. clinical trials.

**Methods:** PBMCs from healthy donors were used to generate knockout T cell models for interferon-γ (IFNγ), Cbl Proto-Oncogene B (CBLB), Fas cell surface death receptor (Fas) and T cell receptor (TCRαβb) genes. The effect of CRISPR-cas9-mediated gene editing on T cells was evaluated using apoptosis assays, cytokine bead arrays and *ex vivo* and *in vitro* stimulation assays.

**Results:** Our results demonstrate that CRISPR/Cas9-mediated gene editing of *ex* vivo T cells is efficient and does not overtly affect T cell viability. Cytokine release and T cell proliferation were not affected in gene edited T cells. Interestingly, memory T cells were more susceptible to CRISPR/Cas9 gene editing than naïve T cells. *Ex vivo* and *in vitro* stimulation with antigens resulted in equivalent antigen-specific T cell responses in gene-edited and untouched control cells; making CRISPR/Cas9-mediated gene editing compatible with clinical antigen-specific T cell activation and expansion assays.

**Conclusion:** Here, we report an optimized protocol for rapid, viable and highly efficient genetic modification in *ex vivo* human antigen specific T cells, for subsequent functional evaluation and/or expansion. Our platform extends CRISPR/Cas9-mediated gene editing for use in gold-standard clinically-used immune-monitoring pipelines and serves as a starting point for development of analogous approaches such as those including transcriptional activators and or epigenetic modifiers.

## INTRODUCTION

CRISPR/Cas9-mediated gene targeting has been used to significantly improve our understanding of T cell biology and discover regulators of e.g. T cell proliferation and differentiation (1–3). To date, most of this work has been performed using cultured peripheral blood mononuclear cells (PBMCs) from healthy donors, or using patient T cells after an initial *ex vivo* expansion (3). In this setting, cell numbers are not limited and there is generally no need to maintain the clonal repertoire of T cells. By contrast, samples obtained within clinical trials in immune oncology are generally limited and antigen-specificity paramount in understanding the underlying immune biology (4–6). Expanding the CRISPR/Cas9 toolbox to antigen-specific T cells from clinical samples would therefore enable further dissection of anti-cancer immune responses and discovery of novel targets for combination immunotherapy (1).

Herein, special consideration should be given to maintaining the *ex vivo* T cells’ phenotype and function during the process of gene engineering (4,7). In addition, a CRISPR/Cas9-mediated gene targeting protocol for clinical samples should ideally be compatible with the timelines used in best-practice immunomonitoring protocols. As such, viral-mediated transfection models, such as lentivirus, adenovirus and retrovirus with variable knockout efficiencies and T cell toxicity are unsuitable (8–10). Ribonucleoprotein (RNP) complexes of Cas9 and a guide RNA (gRNA), when introduced by electroporation, have been reported as suitable for *ex vivo* modification of T cells, but whether a high-level of viability and clonality can be maintained for incorporation into immune monitoring protocols remains unclear (8,9).

Here, we report on an optimized protocol for rapid, viable, and highly efficient genomic manipulation in *ex vivo* human antigen-specific T cells. We demonstrate the efficacy of this gene editing system within common immunomonitoring pipelines and across gRNAs. This system is robust across donors and useable for gene editing in both memory and naïve T cells for studying recall and *de novo* responses, respectively.

## RESULTS

### CRISPR/Cas9-mediated gene editing of ex vivo T cells does not overtly affect T cell viability

We set out to optimize a CRISPR/Cas9-mediated gene editing protocol compatible with PBMC samples analogous to those routinely procured during clinical trials. PBMCs were processed according to best practice operating protocols for PBMC isolation, cryopreservation and cell density used in storage. Using these standardized PBMC aliquots, we first optimized RNP complex electroporation using efficiency and viability as main parameters. As proof-of-concept, we targeted IFNγ as it is not expressed *ex vivo* in T cells, readily induced upon activation and easily assessable by ELISA or intracellular flow cytometry. Electroporation of Cas9/IFNγ-gRNA RNP complexes into T cells resulted in almost complete abrogation of IFNγ expression in both CD8 and CD4 T cells (figure 1a, c). Electroporation, inclusion of Cas9/gRNA complexes and/or crRNA alone, had no effect on T cell viability as measured using amine-reactive dye staining (figure 1b, e). By varying experimental parameters, we observed that while Cas9 protein concentration did not affect T cell viability, the size of the electroporation cuvette was critically important, resulting in a reduced relative viability when using 20 μL cuvettes compared to 100 μL cuvettes (figure 1d). IFNγ knockout efficiency was similar for all experimental conditions and in both CD4 and CD8 T cells (figure 1d). As lower doses of Cas9 protein were equally effective, we chose 100 μg/mL in 100 μL cuvettes as optimal concentration in all subsequent experiments. Analysis of additional parameters of cell viability: 7-AAD, Phosphatidyl Serine (PS) exposure and caspase activation confirmed the minimal loss of cell viability under these conditions (figure 1e). Non-T cells in the PBMC fraction were more sensitive to the T cell optimized protocol, as evident from increased cell death in the total PBMC fraction (figure 1f).

**Figure 1.**
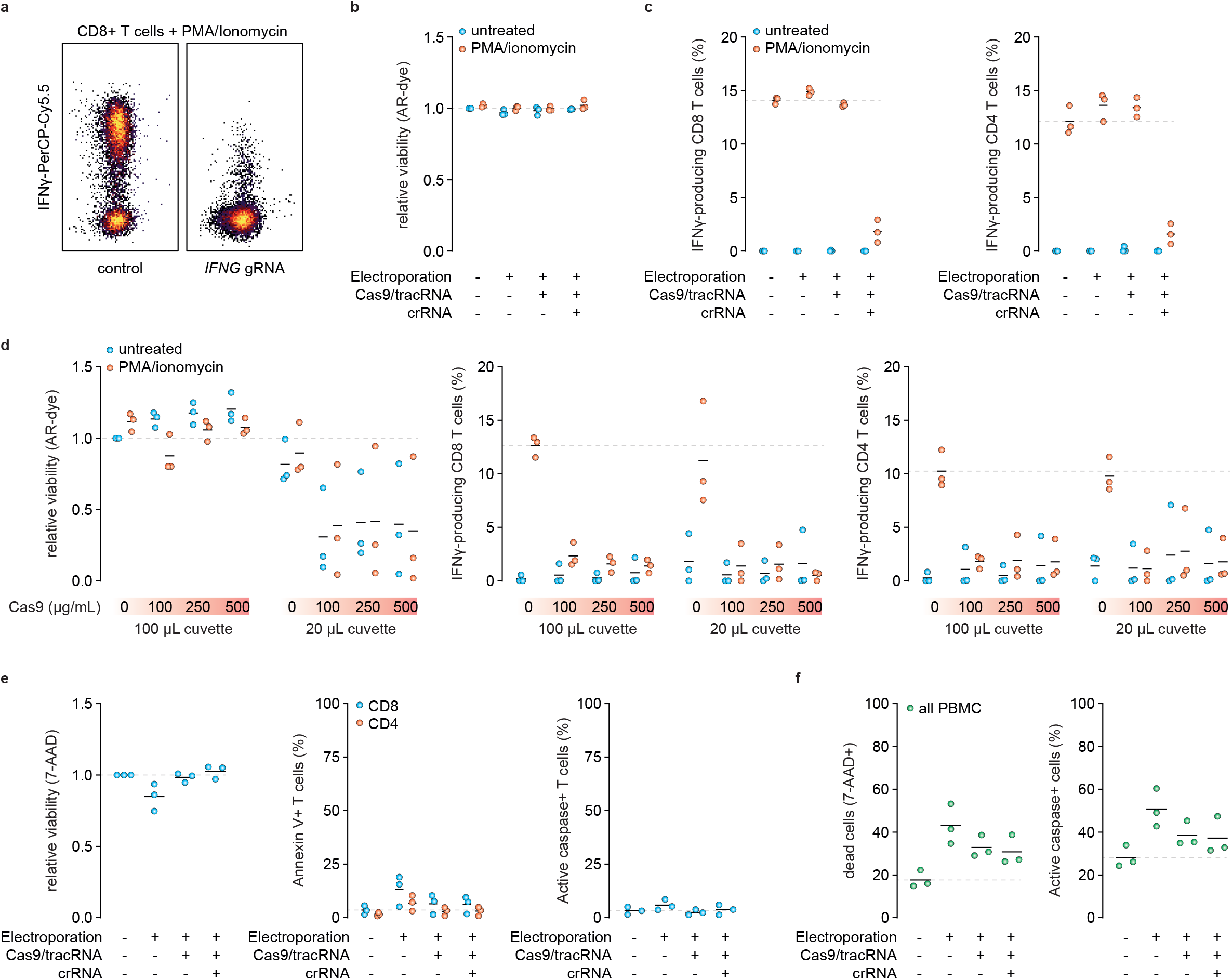
CRISPR/Cas9 electroporation *ex vivo* T cells does not affect T cell viability. Electroporation of PBMCs with Cas9/IFNγ gRNA RNP complexes results in almost complete abrogation of IFNγ expression in both CD8 and CD4 T cells. 16 hours after electroporation, PBMCs were stimulated for 4 hours with PMA/Ionomycin. (A, C) Representative flow cytometry contour plot showing the expression of IFNy production in CD8 T cells. (B) Scatterplots demonstrating the viability of PBMCs after electroporation alone, with Cas9/gRNA complexes and/or crRNA alone. Viability was measured using amine reactive-dye. (D) Effect of different cuvette sizes (100 and 20μL) and different Cas9 protein concentrations (0, 100, 250 en 500 μg/ml) on cell viability. (E) Analysis of additional parameters of cell viability was performed using 7-AAD, Phosphatidyl Serine (PS) exposure and caspase activation in CD8 and CD4 T cells(E) and in the total PBMC fraction(F). Blue dots represent unstimulated PBMCs; orange dots represent PBMCs stimulated with PMA/ionomycin for 4 hours; green dots represent all PBMCs; in e, middle scatterpot, blue dots represent CD8 T cells and orange dots CD4 T cells.

### CRISPR/Cas9-mediated gene editing of ex vivo T cells does not overtly affect T cell function

Next, we tested whether our optimized protocol affected T cell function by evaluating cytokine production and proliferation of gene edited T cells. While release of IFNγ into the extracellular space was markedly reduced in IFNγ knockout T cells, no differences were observed in the release of other cytokines, such as TNFα and IL-2 (figure 2a, b, supplementary figure 1). Atypical induction of IL-4, IL-5 or IL-10 were also not observed under these conditions (figure 2a, b). We also analyzed whether *ex vivo* gene engineered T cells required recovery following electroporation by activating T cells using anti-CD3/CD28 beads at 0, 30, 60, 120 and 240 minutes after RNP nucleofection, followed by 2 day culture and restimulation using PMA/ionomycin. Under all conditions, production of IFNγ, but not TNFα or IL-2, was specifically abrogated (figure 2c, 2d). Accordingly, anti-CD3/CD28 bead- or PHA-based expansion of T cells directly following electroporation did not impair expansion capacity over a period of 14 days (figure 2e). Furthermore, gene-edited T cells that expanded over this period remained IFNγ-negative, suggesting the optimized protocol did not negatively affect T cell fitness when compared with unedited T cells (figure 2f and supplemental figure 1). We reasoned that the direct *ex vivo* gene editing, followed by immediate T cell activation, could even be leveraged to knockout genes critical for anti-CD3/CD28 bead-based activation, as the steady-state protein levels in the absence of transcription are likely sufficient to drive signaling during the initial 24-48 hours of culture. To test this hypothesis, we disrupted expression of the T cell receptor (TCRαβ) complex. T cells were activated using anti-CD3/CD28 beads directly after RNP electroporation. Again, CRISPR Cas9/TCRαβ-gRNA RNP complex electroporation resulted in high TCRαβ knockout efficiency in both CD8 and CD4 T cells at 48h (figure 2g). As anticipated, TCRαβ knockout T cells proliferated comparably to non-edited T cells over a period of 14 days (figure 2h). As for IFNγ, no detrimal effects of gene editing were observed and near-complete loss of TCRαβ was maintained at day 14 (figure 2i).

**Figure 2.**
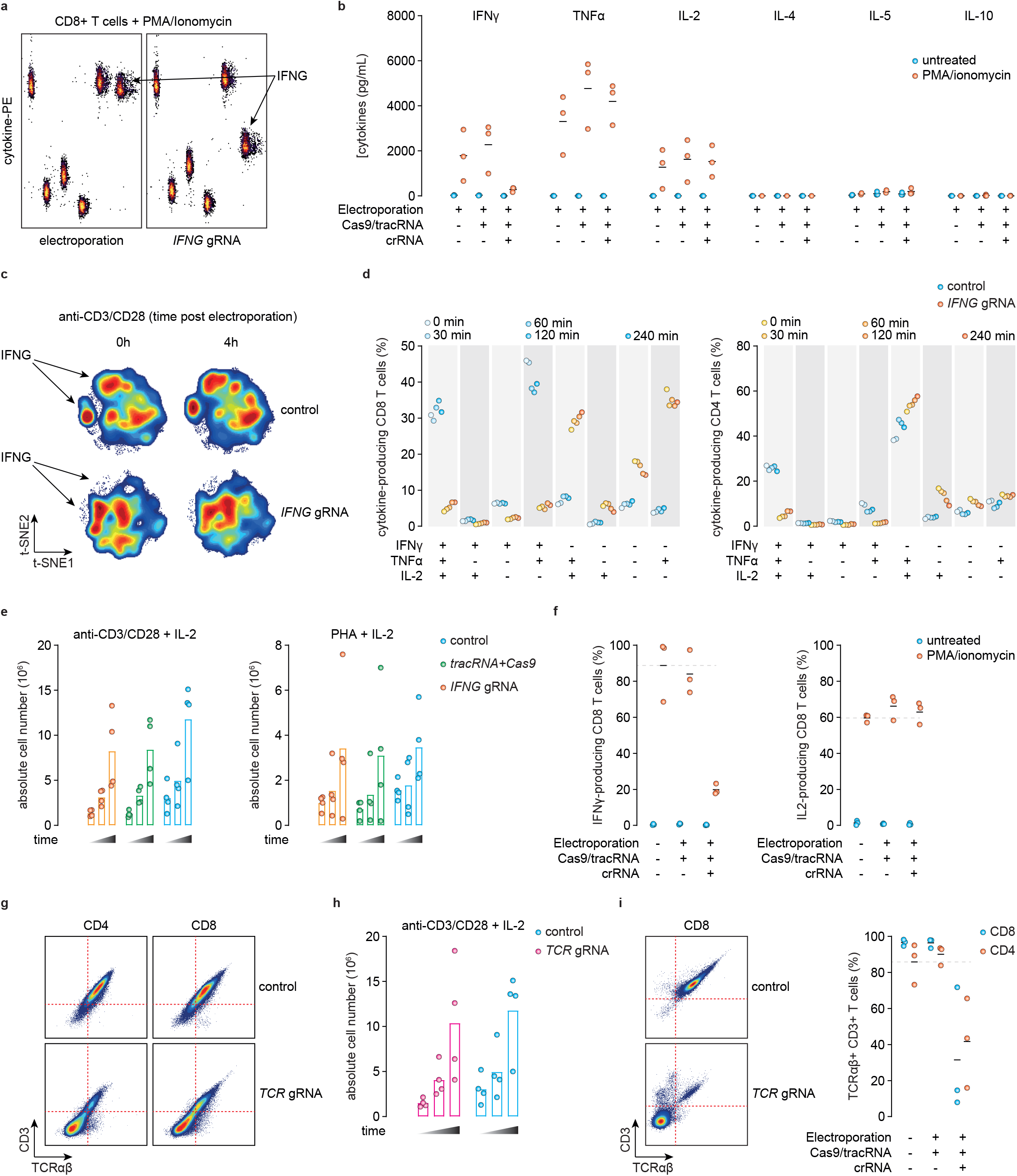
CRISPR/Cas9-mediated gene editing of *ex vivo* T cells does not affect T cell function. IFNγ, TNFα, IL-2, IL-4, IL-5 and IL-10 release was evaluated using a cytokine bead array kit. (A,B) Electroporation with Cas9/IFNγ gRNA RNP complexes results in reduced IFNγ release whereas no differences are observed in the release of TNFα and IL-2, 2 days after electroporation. Atypical induction of IL-4, IL-5 or IL-10 are also not observed. (C) t-SNE plots showing the reduced IFNy release after stimulation with beads at t=0 and t=4 hours after electroporation. (D) IFNγ, TNFα and IL-2 cytokine-producing CD8 T cells after activation with anti-CD3/CD28 beads at 0, 30, 60, 120 and 240 minutes after RNP nucleofection, followed by 2 day culture and restimulation using PMA/ionomycin. (E) Scatterplot showing the anti-CD3/CD28 bead- or PHA-based expansion of T cells directly following electroporation over 14 days. (F) IFNγ and IL-2 production of 14 days expanded gene-edited CD8 T cells. (G) Flow cytometry contour plot showing the TCRαβ knockout in CD8 and CD4 T cells 48 hours after electroporation (G). (H) Expansion capacity of anti-CD3/CD28 activated TCRαβ knockout cells over 14 days. (I) Flow cytometry contour plot left) and scatterplot (right) showing the TCRαβ knockout in CD8 and CD4 T cells after 14 days.

### CRISPR/Cas9-mediated gene editing of ex vivo T cells preferentially targets memory T cells

We next determined whether memory or naïve T cells were more amenable to CRISPR/Cas9-mediated gene editing in the *ex vivo* setting. PBMCs were subjected to TCRαβ-targeting RNP electroporation, followed by 8-day culture with cytokines IL-2, IL-15 and/or IL-7 to support memory and naïve T cells, respectively. Cell viability was similar across all conditions (figure 3a). Percentages of memory CD8 T cells were similar between TCRαβ knockout and TCRαβ-expressing cells regardless of the cytokine cocktail used (figure 3b,c). Notably, there was a slight increase in the percentage of memory CD4 T cells after TCRαβ knockout compared to the TCRαβ-proficient cells, independent of cytokine stimulation (figure 3c).

**Figure 3.**
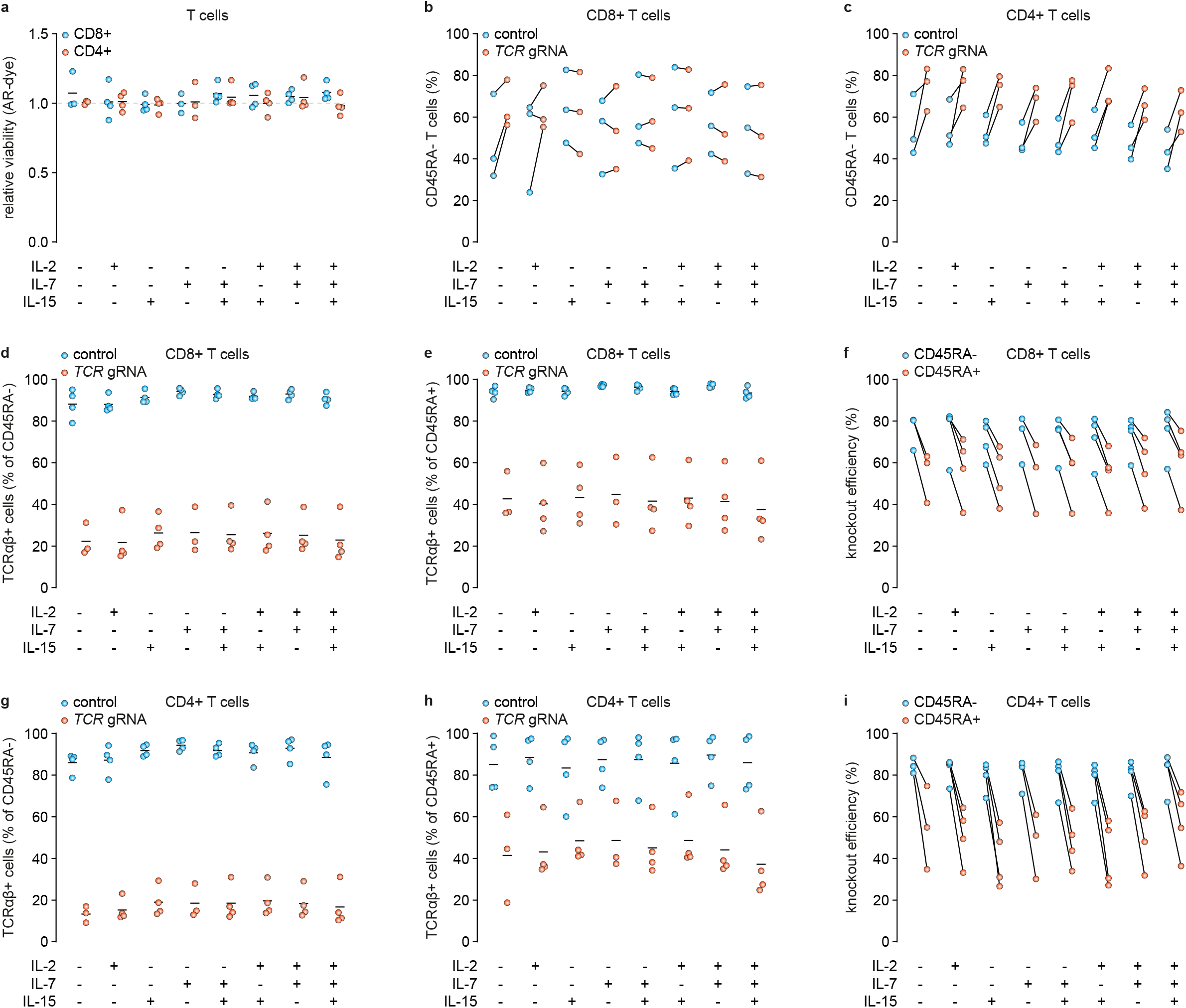
CRISPR/Cas9 gene editing preferentially targets memory T cells. Gene editing of TCRαβ, IFNγ, CBLB or FAS in PBMCs. (A) Viability of CD8 versus CD4 TCRαβ knockout T cells after *in vitro* stimulation with IL-2, IL-15 and/or IL-7 for eight days. (B) Percentages of CD8 and (C) CD4 memory T cells within TCRαβ knockout T cells compared to TCRαβ-proficient cells after co-culturing with IL-2, IL-15 and/or IL-7 for eight days. (D) Percentages of CD8 memory and (E) CD8 naïve T cells expressing TCRαβ eight days after gene editing. (F) Gene editing efficiency of CD8 memory T cells compared to CD8 naïve T cells. (G) Percentages of CD4 memory and (H) CD4 naïve T cells expressing TCRαβ eight days after gene editing of the TCRαβ receptor. (I) Gene editing efficiency of CD4 memory T cells compared to CD4 naïve T cells. Culture conditions as indicated.

To further explore this observation, we investigated the knockout efficiency within the memory and naïve T cell populations (figure 3d-i). The knockout efficiency of memory T cells varied between 60-85%, in CD8^+^ T cells and between 70-95% in CD4^+^ T cells (figure 3d,g). By contrast, the knockout efficiency for naïve T cells ranged between 35-80% in CD8 T cells and between 30-80% in CD4 T cells (figure 3e,h). In addition, the percentage of naïve TCRαβ knockout CD8 T cells was higher when compared to naïve TCRαβ knockout CD4 T cells (figure 3e,h). These findings collectively suggest that memory T cells, especially memory CD4 T cells, are slightly more susceptible to CRISPR/Cas9 gene editing than naïve T cells.

### CRISPR/Cas9-mediated gene editing is compatible with clinical antigen-specific T cell activation and expansion assays

Finally, having established the high viability and efficacy of our approach, we evaluated whether our optimized protocol was compatible with best-practice assays for monitoring immune responses in clinical trial settings. We analyzed *ex vivo* peptide stimulation, as well as *in vitro* stimulation (IVS) assays. Using CEF peptides as model antigen, and T cell activation markers CD69 and CD137 as a readout, we were able to demonstrate equivalent CEF-induced responses in gene-edited and untouched control cells for both the *ex vivo* assay (figure 4a, c) and following IVS (figure 4b, d). These responses were observed for IFNγ knockouts, as well as for Cbl Proto-Oncogene B (CBLB) and Fas cell surface death receptor (Fas) gRNAs previously reported by others (figure 4a-d).

**Figure 4.**
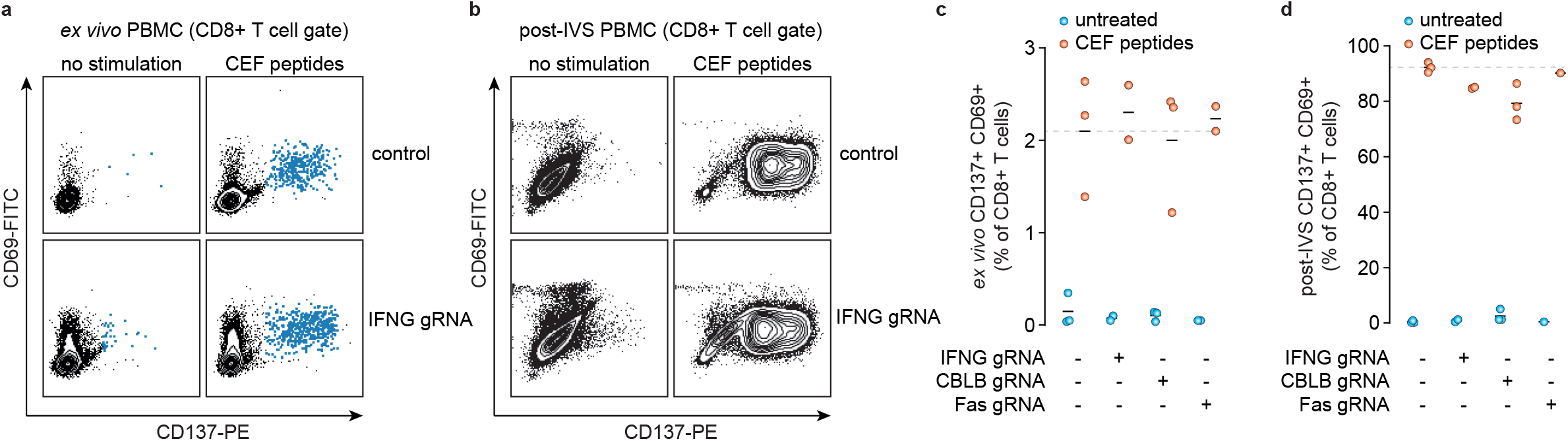
CRISPR/Cas9 editing does not influence the antigen-specificity of T cells. Gene editing of IFNγ, CBLB or FAS in PBMCs. (A) Representative flow cytometry contour plots showing the expression of T cell activation markers CD69 and CD137 after *ex vivo* and (B) post-IVS stimulation with CEF. (C) Scatterplot representing the percentage of CD8 T cells expressing both activation markers CD137 and CD69 within variable CRISPR gene editing conditions and untouched control cells after *ex vivo* stimulation at day 2 and (D) post-IVS stimulation with and without CEF peptides at day 13. Blue dots represent PBMCs not stimulated with CEF peptides and orange dots PBMCs stimulated with CEF.

## DISCUSSION

We describe a rapid, robust and flexible platform for CRISPR/Cas9-mediated gene editing in primary *ex vivo* human T cells. We have optimized efficiency, viability, as well as culture conditions to allow rapid and direct *ex vivo* modification of T cells for subsequent functional evaluation and/or expansion. Our platform is amenable to integration into clinically used pipelines for immune response evaluation using antigens of choice.

CRISPR/Cas9-mediated gene editing has become an attractive approach for modifying T cells, due to its simplicity, operability, low costs and capability of multiplex genome editing (9,11). However, to date, limited experience on the application of CRISPR/Cas9 electroporation in immune monitoring assays in clinical studies is available. The prerequisites for manipulating T cells in immunoassays are maintaining viability, antigen-specificity and function (4,7). In this study, we demonstrate for the first time that these prerequisites are not affected by CRISPR/Cas9 gene editing, thereby providing a proof-of-principle for the application of CRISPR/Cas9 in immune monitoring assays. In both CD8 and CD4 T cells, a highly efficient gene knock-out up to 90% was reached after a single transfection. Previously published literature, using CRISPR/cas9 RNP electroporation, reported efficiency rates of 20-90% with low efficiency rates in resting primary T cells (9,12–16). Moreover, we demonstrated that CRISPR/Cas9-edited cells could be maintained without stimulation in culture for up to 7 days, or supplemented with a diverse range of cytokines without comprising gene knockout efficacy. Activation of T cells directly, or at a later time-point of choosing did not negatively affect T cell function, expansion, nor did cryopreservation and subsequent restimulation. Finally, while the current work focused on knockout of genes, recent advances in non-homologous end joining (NHEJ)-mediated template incorporating using CRISPR/cas9 should be amenable to the current platform, allowing more complex *ex vivo* T cell engineering (12,14,17).

Advances in descriptive immune monitoring have expanded our understanding of anti-cancer immune responses. High-parameter flow, mass and spectral cytometry have allowed the simultaneous assessment of most immune cell populations *ex vivo* (18,19). Imaging using e.g. PET/SPECT allows non-invasive assessment of specific immune cells or immune checkpoints throughout the body, and transcriptomic approaches provide ever deeper dimensional assessment of gene expression, including spatial organization (20–22). By contrast, *ex vivo* functional assays have remained largely unchanged over the years, with peptide stimulation and intracellular flow cytometry, ELISPOT or ELISA used for functional readout on one or two hallmark activation markers (4,23). The platform we present here significantly expands the potential of these assays. By seamlessly integrating CRISPR/Cas9-mediated gene editing into *ex vivo* immune monitoring pipelines, it is possible to assess how T cells from clinically-treated patients differentially depend on genes-of-interest, but also screen *ex vivo* for novel combination targets to further augment T cell activation (14,15). As demonstrated here, these targets need not be limited to cell surface targets accessible by antibody-based therapeutics (14,15). One interesting observation is that naïve CD4, but not CD8, T cells appear less sensitive to CRISPR/Cas9-mediated gene editing than their memory counterparts. Whether this is the result of the optimized electroporation protocol used, or an intrinsic feature of naïve CD4 T cells is currently unknown. Nevertheless, targeted genes of interest could be knocked out in ∼60% of CD4 T cells, establishing our platform as an effective approach for studying both recall and *de novo* immune responses. An interesting approach would be to further combine our platform with recent advances in CRISPR/Cas9-mediated gene editing in monocyte-derived dendritic cells. This could allow reciprocal knockout of ligand-receptor interactions of particular interest for complex ligand-receptor pairs, such as CTLA-4/CD28 and CD80/CD86.

Taken together, our platform represents an advance on previous reports on CRISPR/Cas9-mediated gene editing and extends this approach to use in gold-standard clinically-used immune-monitoring pipelines.

## MATRIALS AND METHODS

### Peripheral Blood Mononuclear Cells

Peripheral blood mononuclear cells (PBMCs) were isolated from blood of healthy volunteers after written informed consent was obtained (Sanquin). For isolation, buffy coats were mixed at a 1:2 ratio with RPMI supplemented with 2,5% FCS, layered on a Ficoll-Paque gradient at a 1:2 volume ratio and centrifuged at 900 g for 20 minutes without brake. PBMC on the interface of the Ficoll-Paque and plasma layers were isolated by pipette (5-10 mL), supplemented with ice-cold PBS to a final volume of 50 mL and centrifuged at 560 g for 8 minutes without brake. PBMC pellets were pooled, supplemented with ice-cold PBS to a final volume of 50 mL and centrifuged at 350 g for 8 minutes without brake. After PBMCs were resuspended in 50 mL of ice-cold PBS, PBMCs were counted using the Bürker counting chamber and centrifuged at 350 g at 8 minutes without brake. Finally, PBMCs were resuspended in 1 mL of freezing medium, consisting of 90% FCS and 10% DMSO, at a concentration of 10-100^6^ cells per cryovial.

### Cas9, tracRNA and crRNA

Cas9 (Alt-R Cas 9 Nuclease V3), tracRNA (Alt-R CRISPR-Cas9 tracRNA) and the crRNAs (Alt-R CRISPR-Cas9 crRNA) were synthesized by Integrated DNA Technologies (Coralville, USA). Each crRNA was reconstituted in nuclease free duplex buffer (Lonza) to a final concentration of 100 μM. See Table 1 for gRNAs and sequences. For TCRαß: equal volumes of TRAC1 and TRBC were mixed before adding the RNP complex to the PBMCs, see RNP electroporation.

### RNP electroporation

PBMCs were counted and resuspended in X-VIVO 15™ at densities as indicated. After optimization as indicated, the final protocol to prepare Cas9/gRNA RNP complexes was as follows: 150 pmol annealed gRNA per nucleofection condition was prepared by combining 1.5 μL tracRNA with 1.5 μL crRNA. The mix was annealed by heating at 95°C for 5 minutes and slowly cooled down to room temperature for 10 minutes. Next, 12 μL duplex buffer (Lonza, Basel, Switzerland) was added to get a final concentration 150 pmol gRNA in 15 μl. To prepare a Cas9/gRNA RNP complex, 10 μg Cas9 (10 μg/ml) was added to the gRNA complex. This mixture was incubated for 10 minutes at room temperature. Finally, duplex buffer was added to the RNP complex to obtain a final volume of 20 μL.

For electroporation, PBMCs were centrifuged and washed once with PBS. Cells were resuspended in 100 μL nucleofection solution at densities as indicated. Nucleofection solution consisted of 82 μL P2 Primary Cell Nucleofector™ Solution (Lonza) and 18 μL Supplement 1 (Lonza, Basel, Switzerland) for 100 μL cuvettes and 16.4 μL P2 Primary Cell Nucleofector™ Solution and 3.6 μL Supplement 1 for 20 μL cuvettes. RNP complex was added to the PBMCs and incubated for 2 minutes at room temperature. The PBMC/RNP mixture was transferred to a nucleofection cuvette, followed by electroporation of cells using a 4D-Nucleofector machine (Amaxa, program P2, pulse code EH100). PBMCs that did not undergo electroporation served as control and were centrifuged and washed with PBS.

### Intracellular cytokine staining

16 hours after electroporation, PBMCs were stimulated for 4 hours with PMA/Ionomycin (eBioscience stimulation cocktail) at 37° degrees. For intracellular staining of IFNγ, tumor necrosis factor alpha (TNFα), and interleukin-2 (IL-2), secretion of cytokines was blocked using golgiplug (protein transport inhibitor; BD 555029). After 4 hours, cells were stained with Zombie Aqua viability dye according to manufacturer’s protocol and washed with PBS supplemented with 5% FCS. Next, cells were fixed and permeabilized (CEL FIXATIE & PERMEABILISATIE KIT, GAS-002, Nordic-Mubio) and stained with surface markers CD3, CD8, CD4, and intracellular markers IFNγ and IL-2 for flow cytometer analysis. For the TCRαβ knockout experiments, only cell surface staining with TCRαβ, CD3, CD8 and CD4 antibodies was performed. See supplementary table 2 for characteristics of the antibodies. For fixation and permeabilization cells were incubated with 100 μL/well of reagent A (fix) for 15 minutes at room temperature. Next, cells were washed with PBS and incubated with a mixture of reagent B (perm) and antibodies for 15 minutes at room temperature. In total 100 μL of the mixture reagent B plus antibodies was added per well. After incubation, cells were washed twice, centrifuged and resuspended in flow cytometry buffer (PBS + 2% FCS).

Intracellular staining of IFNγ, TNFα and IL-2 was also performed on engineered T cells activated with anti-CD3/CD28 beads (Thermofisher, Cat. 11131D) at 0, 30, 60, 120 and 240 minutes after RNP nucleofection, followed by 2 day culture and restimulation using PMA/ionomycin.

### Apoptosis assay

Apoptosis was assessed 24 hours after CRISPR/Cas9-mediated gene editing. PBMCs were counted, centrifuged and resuspended in X-VIVO 15™ at a density of 1-2×10^6^ cells/mL. Subsequently, 1 μL Violet live Caspase probe per 0.5 mL of cell suspension was added and cells were incubated for 45 minutes at 37°C. After washing, cells were resuspended in fresh medium and incubated for an additional 30 minutes at 37°C while protected from light. Afterwards, CD8 and CD3 antibodies were added and cells were incubated for 30 minutes at room temperature. Cells were washed twice with cold PBS and resuspended in 1x Annexin V Binding Buffer at a final concentration of 1×10^6^ cells/mL. 1×10^5^ cells were incubated with 5 μL of Annexin V and 5 μL of 7-Aminoactinomycin D Invitrogen™ (7-AAD) for 15 minutes at room temperature in the dark. Cells were analyzed within 1 hour using the BD FACSVerse flow cytometer (BD Biosciences).

### Cytokine bead array

One day after electroporation (10×10^6^ cells in 2 mL, 1 mL per well), gene-edited and control PBMCs were incubated with PMA/ionomycin (eBioscience stimulation cocktail) for 4 hours at 37C°. Subsequently, supernatant was collected, and stored at −80°C.

The human Th1/Th2 Cytokine Assay (BD™ Cytometric Bead Array (CBA) Human Th1/Th2 Cytokine Kit II) was used to quantitatively measure cytokine production according to the manufacturer’s protocol. In brief, supernatant was thawed and 50 μL was used for the cytokine bead array. Samples and standards were incubated with the capture beads for 3 hours at room temperature, while protected from light (Supplementary data, supplementary table 2). After incubation, assay tubes were washed, centrifuged and resuspended before flow cytometer analysis.

### T cell expansion assay

PBMCs were plated at a density of 5×10^6^/ml in a 24 wells plate. Expansion of gene-edited PBMCs was tested by stimulation with CD3/CD28 beads (20 μL per well, 5×10^6^ cells per well) or phytohemagglutinin (L1668 Sigma)(PHA, 10 μg/mL). IL-2 (50 IU/mL) was added to both stimulating conditions. Stimulation was done directly after electroporation. Plates were incubated at 37°C and cells were counted and at day 7, 10 and 14. Cell density was maintained around 1×10^6^ cells/mL and medium, IL-2 and/or PHA was refreshed at day 7, and 10.

### *Ex vivo* T cell stimulation

PBMCs were resuspended in 500 μL culture medium (X-VIVO 15™ with 10% human serum) and plated in 100 μL/well of 96-well round bottom plates (Thermofisher) at a density of 1×10^6^/ml. Cells were *ex vivo* stimulated either with viral peptides consisting of a pool of 23 different peptides originating from Cytomegalovirus, Epstein-Barr virus and influenza virus peptides (CEF) or with interleukin 7 (IL-7) and interleukin 15 (IL-15). CEF was synthesized by JPT Peptide Technology (Berlin, Germany). Recombinant human IL-7 (Asp26-His177, size 10 μg) and recombinant human IL-15 (amino acids Asn49-Ser162, size 10 μg) were purchased from Biolegend.

For *ex vivo* stimulation with CEF, cells were activated one day after electroporation by incubation with CEF peptides at a concentration of 1 μg/ml at 37°C. For *ex vivo* stimulation with IL-7 and IL-15, cells were activated by incubation with IL-7 and IL-15 at a concentration of 25 ng/ml at 37°C at day 8. After 16-26 hours of incubation with CEF or IL-7 and IL-15, cells were stained for T cell phenotype with CD3, CD8 and CD14 and for T cell activation with CD69 and CD137, see supplementary table 2.

### *In vitro* T cell stimulation (IVS)

PBMCs were plated in 2 mL culture medium per well per CRISPR condition of 24-wells plates at a density of 5×10^6^/ml. Culture medium composed of X-VIVO 15™ enriched with 10% human serum, 1% penicillin/streptomycin (P/S), 1% L-glutamine (L-glu) and IL-2 (20 IU/ml). PBMCs were stimulated with CEF for 13 days. A concentration of 1 μg/mL CEF was used. Medium was changed at day 4, 6, 8 and 11, according to the harmonization protocol of short-term *in vitro* culture for expansion of antigen-specific CD8+ T cells (4). At day 13, cells were re-stimulated with CEF at a concentration of 1 μg/mL overnight at 37°C and 5% CO_2_, while medium served as a negative control and PHA (L1668 Sigma) at 1 μg/mL as positive control. After 16-26 hours cells were stained for phenotype with CD3 and CD8 and for activation with CD69 and CD137, see supplementary table 2 for characteristics of the antibodies.

### Memory T cells assay

PBMCs were resuspended in 800 μL culture medium (X-VIVO 15™ with 10% human serum) and plated in 100 μL/well of 96-well round bottom plates (Thermofisher) at a density of 0,625×10^6^/mL. PBMCs were stimulated with IL-2, IL-7 and/or IL-15 at a concentration of 25 ng/mL or left untreated as indicated. Cells were cultured for 7 days. At day 8, cells were stained for T cell memory phenotype with CD4, CD8, CD45 and TCRα/ß, see supplementary table 2 for characteristics of the antibodies.

## SUPPLEMENTARY TABLES

**Supplementary table 1.**
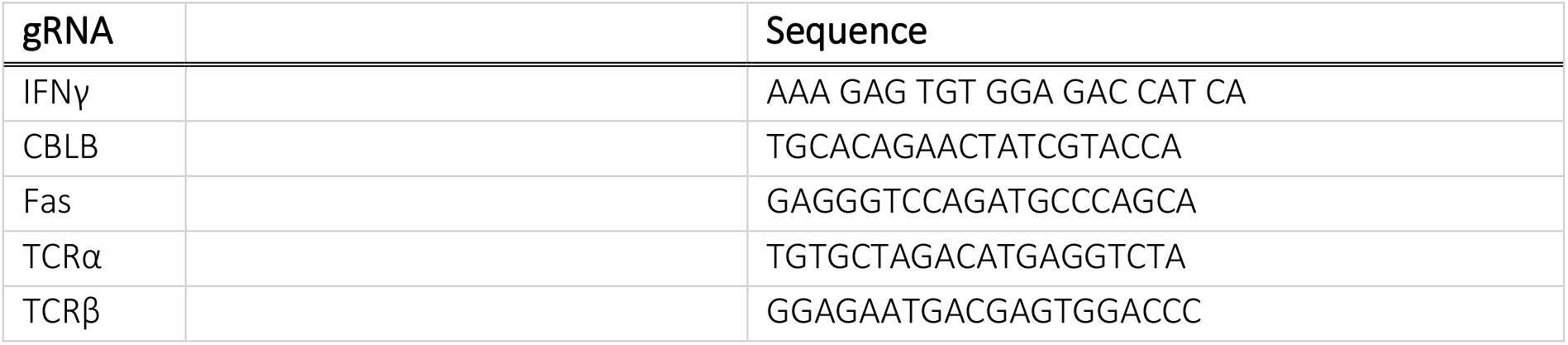

**Supplementary table 2.**

| Antibody | Color | Cat No | Supplier | Experiment |
| --- | --- | --- | --- | --- |
| CD3 | APC-eFluor 780 | 47-0038-42 | eBioscience™,<br>ThermoFisher | *Ex vivo* and *in vitro* T cell stimulation |
| CD3 | PE | 12-0038-41/42 | eBioscience™,<br>ThermoFisher | Apoptosis assay |
| CD3 | FITC | 555332 | BD | Intracellular cytokine staining |
| CD4 | PE | 12-0048-42 | eBioscience™,<br>ThermoFisher | Intracellular cytokine staining<br>Memory T cell assay |
| CD8 | BV421 | 562428 | eBioscience™,<br>ThermoFisher | Intracellular cytokine staining<br>*Ex vivo* and *in vitro* T cell stimulation<br>Memory T cell assay |
| CD8 | APC-eFluor 780 | 47-0088-42 | eBioscience™,<br>ThermoFisher | Apoptosis assay<br>Intracellular cytokine staining |
| CD45RA | Pe-Cy7 | 25-0458-41 | eBioscience™,<br>ThermoFisher | Memory T cell assay |
| CD69 | FITC | 11-0699-41 | eBioscience™,<br>ThermoFisher | *Ex vivo* and *in vitro* T cell stimulation |
| CD137 | PE | 12-1379-42 | eBioscience™,<br>ThermoFisher | *Ex vivo* and *in vitro* T cell stimulation |
| TCR$\alpha/\beta$ | APC | 17-9986-41 | eBioscience™,<br>ThermoFisher | Memory T cell assay<br>Intracellular cytokine staining |
| IFNγ | PerCP-Cy5.5 | 45-7319-42 | eBioscience™,<br>ThermoFisher | Intracellular cytokine staining |
| TNFα | APC | 17-7349-82 | eBioscience™,<br>ThermoFisher | Intracellular cytokine staining |
| IL-2 | PE-Cy7 | 25-7029-42 | eBioscience™,<br>ThermoFisher | Intracellular cytokine staining |

**Supplementary table 3.**

| Tube label | Concentration (pg/mL) | Cytokine Standard dilution |
| --- | --- | --- |
| 1 | 0 (negative control) | No standard dilution (Assay Diluent only) |
| 2 | 20 | 1:256 |
| 3 | 40 | 1:128 |
| 4 | 80 | 1:64 |
| 5 | 156 | 1:32 |
| 6 | 312,5 | 1:16 |
| 7 | 625 | 1:8 |
| 8 | 1250 | 1:4 |
| 9 | 2500 | 1:2 |
| 10 | 5000 | Top Standard |

## FIGURE DESCRIPTIONS

**Supplementary figure 1. CRISPR/Cas9-mediated gene editing of *ex vivo* T cells does not affect T cell function.**

Scatterplots representing the IFNy-, IL-2, and TNFα producing CD8 and CD4 T cells after activation with anti-CD3/CD28 beads, or PHA, 14 days after expansion (relates to figure 2f).

**Supplementary figure 2. Gating strategy.**

Scatterplots representative for the gating strategy used throughout the manuscript. The depicted scatterplots show gene edited PBMCs after CBLB knockout and IVS stimulation with CEF. Cell populations are depicted above the FACS images and the gated population are depicted below.

